# Why development time matters: temperature influences plasticity while nutritional stress reshapes selection and evolutionary potential

**DOI:** 10.64898/2026.10.31.763280

**Authors:** Greg M. Walter, Avishikta Chakraborty, Fiona Cockerell, Lesley A. Alton, Craig R. White, Matthew D. Hall, Carla Sgrò

## Abstract

The time it takes organisms to reach adulthood is often considered a central life-history trait that influences fitness and is highly plastic across environments. Despite its importance, direct estimates of how selection on development time varies across combinations of environmental stressors remain rare, limiting our understanding of its role in population responses to environmental change. We reared 68 full-sibling families of the vinegar fly, *Drosophila melanogaster*, at two temperatures (14°C and 28°C) on two diets (standard and low-calorie), and measured development time, body size (mass and wing size), egg-to-adult viability and productivity (number of offspring). Temperature caused the largest plastic response, with development taking 4–5 times longer at 14°C, whereas the low-calorie diet extended development by 20–30%. Yet diet, not temperature, changed directional selection on development time. On the standard diet, faster development was associated with higher fitness at both temperatures. On the low-calorie diet, this advantage disappeared, and at 14°C slower-developing females tended to produce more offspring. Viability selection showed consistent patterns. The low-calorie diet also increased evolvability in development time c.10–16-fold at both temperatures, and genetic correlations between diets were close to zero, indicating strong genotype-by-environment interactions. Plasticity in development time was not associated with fitness in any treatment. Overall, we provide evidence that development time is a functional trait whose relationship with fitness depends on the environment. Nutritional stress induced a smaller plastic response in development time than temperature, but had a larger effect on directional selection and evolvability. The stressor causing the greatest plastic change in a functional trait is therefore not necessarily the one that shapes its selection and evolution.

## Introduction

Ongoing global change is threatening natural populations with warmer and more unpredictable thermal environments (Parmesan & Yohe 2003; Parmesan 2006). Shifting thermal regimes then shape changes in the biotic environment, which can influence the abundance and distribution of food resources (Cross *et al*. 2015; Hardison & Eliason 2024). Such multidimensional environmental change alters how organisms interact with their physical environment, threatening population persistence if individuals struggle to survive and reproduce under the new conditions (Buckley *et al*. 2017). Despite the need to understand population resilience to realistic climate change scenarios, experiments rarely test how combinations of stressors reshape selective pressures acting on ecologically important traits (Kokko *et al*. 2017; Siepielski *et al*. 2017).

Plastic changes in phenotype that occur within a generation allow organisms to shift life history traits to track environmental change (Nylin & Gotthard 1998; Acasuso-Rivero *et al*. 2019; Smith *et al*. 2025). However, such plasticity has limits that are likely to be exceeded under novel or extreme conditions, which creates fitness costs that threaten population persistence (DeWitt, Sih & Wilson 1998; Chevin, Lande & Mace 2010; Hoffmann & Bridle 2023). When the limits to plasticity are reached, population persistence increasingly depends on evolutionary responses that require selection on heritable variation (Gomulkiewicz & Holt 1995; Hoffmann & Sgrò 2011). Fitness costs associated with environmentally induced (plastic) phenotypic change can change the direction or strength of selection on traits important for survival and reproduction, yet such effects are rarely quantified (Kingsolver *et al*. 2012; Siepielski *et al*. 2017; Westneat *et al*. 2019). Assessing population persistence under global change therefore requires understanding how plasticity and selection respond to realistic combinations of environmental stressors, and how their interaction is likely to shape evolutionary responses (Gienapp *et al*. 2008; Chevin, Lande & Mace 2010; Urban *et al*. 2024).

The time it takes organisms to reach adulthood is a functional life history trait that shapes adult body size, reproductive timing, predator escape and competitive ability, and is therefore expected to be under strong selection (Nylin & Gotthard 1998; Monaghan 2008; Kasumovic *et al*. 2013). Despite its ecological importance, there are surprisingly few direct estimates of natural selection acting on development time, particularly as environments increase in stress. Reciprocal transplant experiments with plants have shown that selection on development time changes with environments, which creates fitness costs in novel environments (Walter *et al*. 2018; Hamann, Wadgymar & Anderson 2021; Pålsson *et al*. 2024; Walter *et al*. 2026c). By contrast, evidence that development time influences fitness in animals is primarily from artificial selection experiments. Selection for faster development reduces larval survival and adult fecundity, revealing fitness costs of shorter development time under control conditions (Gotthard, Nylin & Wiklund 1994; Zwaan, Bijlsma & Hoekstra 1995; Chippindale *et al*. 1997; Prasad & Joshi 2003; Yadav & Sharma 2014). However, artificial selection experiments do not estimate how natural selection acts on individuals across changing environments, and so provide limited insight into whether the direction or strength of selection on development time vary across ecological contexts (Zwaan, Bijlsma & Hoekstra 1995; Kingsolver *et al*. 2001). Manipulative experiments are therefore needed to test how genotypes from a natural population change their development time to match the environment, and the consequences for fitness (Metcalfe & Monaghan 2003).

Because plasticity changes the phenotypes that selection acts on, predicting evolutionary responses to environmental change requires understanding how selection varies across environments, and how genetic variation is expressed under those same conditions (Ghalambor *et al*. 2007; Gienapp *et al*. 2008; Chevin, Lande & Mace 2010). Where genotypes vary in their fitness or phenotypic response to environmental change, genotype-by-environment interactions (G×E) emerge. Genotypes can vary in the strength of their response to changes in the environment, which alters the amount of genetic variation expressed, and/or genotypes can respond differently to environmental change, which weakens cross-environment genetic correlations to <1 (Falconer 1952; Via & Lande 1985). G×E can therefore determine the amount of genetic variation expressed within a population as well as whether different genotypes are favoured as environments change. As a consequence, genetic variance can either increase under stressful conditions if stress exposes genetic differences, or it can decrease if stress erodes genetic variation or inflates environmental variance (Hoffmann & Merilä 1999; Charmantier & Garant 2005). Where selection and genetic variation change with the environment, the evolutionary potential of populations to respond to global change will be substantially altered (Hoffmann & Sgrò 2011).

Combinations of environmental stressors can activate physiological and developmental pathways that differ from those induced by single stressors (Crain, Kroeker & Halpern 2008; Todgham & Stillman 2013; Cross *et al*. 2015). In natural populations, warmer temperatures frequently coincide with reduced or less predictable food availability, raising the possibility that selection on key life-history traits may be unstable or even conflicting across environments (Holden *et al*. 2019; King *et al*. 2022). Under such scenarios, adaptive responses may be slowed, or alternative life history strategies favoured (Botero *et al*. 2015). Incorporating nutritional stress alongside temperature is therefore important for understanding evolutionary and demographic responses to climate change (Rosenblatt & Schmitz 2016). Populations exposed to warmer environments or higher densities typically develop faster, while diet restriction and cooler temperatures slow development (Stearns & Koella 1986; Prasad & Joshi 2003; Trotta *et al*. 2006). Whether such plastic responses are adaptive, and whether selection and genetic variation for development time shift across combinations of stressors, remains poorly understood.

Here, we test how combinations of temperature and nutritional stress shape selection, plasticity and evolutionary potential of development time while accounting for body size. Using 68 full-sibling families from a large mass-bred population of the vinegar fly, *Drosophila melanogaster*, we exposed larvae from each family to two developmental temperatures (14°C and 28°C) and two diets (standard and low-calorie 25% dilution), and measured development time, body size, and productivity (our fitness proxy) on individual females, as well as egg-to-adult viability for each family. This approach allowed us to evaluate not only whether development time is under selection, but how environmental stressors alter both the selective landscape and the genetic variation that determines evolutionary potential under global change. We distinguish three ways in which development time and fitness can be linked. First, an environment can change mean development time (plasticity). Second, within a given environment, individuals or genotypes that develop faster or slower than others can differ in fitness (selection on development time). Third, genotypes can differ in how much their development time changes between environments, which can itself be associated with fitness (selection on plasticity). These may not align: an environment can lengthen development for all genotypes while selection within that environment still favours those that develop fastest. We predicted that combinations of temperature and diet would together shape selection and the expression of genetic variance in development time, which would influence evolutionary potential.

## Materials and Methods

We constructed a large mass-bred population of *D. melanogaster* by combining populations from eight sampled locations from a c.300×600 km area across Victoria, Australia (Walter *et al*. 2026a). As described in Walter *et al*. (2026b), we combined 500 individuals (250 virgin females and 250 males) from each of the eight populations across 20 bottles (c.500 flies per bottle), which we maintained at 25°C for three generations to allow thorough mixing. Combining populations increased the phenotypic and genetic variation available for estimating selection, and represents genetic variation in a larger population connected by gene flow (Kennington, Gockel & Partridge 2003). We then used the mass-bred population in two experiments. In experiment 1, we generated 750 full-sibling families and measured their development time in a cold, low-calorie treatment. In experiment 2, we used families from the tails of the development time distribution to estimate selection on development time and body size at two temperatures and on two diets.

### Experiment 1: Establishing the full-sibling families

From the large population, we established full-sibling families using a nested half-sibling breeding design (Lynch & Walsh 1998), where we mated 150 sires each to five females, generating 750 full-sibling families. Walter *et al*. (2026b) used this breeding design to estimate additive genetic variance in fitness across six combinations of temperature (14°C, 25°C and 28°C) and diet (standard and low-calorie). The low-calorie diet was a 25% dilution representing reduced food availability predicted under climate change (Chakraborty *et al*. 2023). Vials containing a single full-sibling family were randomised, and eclosing adults were scored daily for development time in the 14°C low-calorie treatment, which showed the greatest variation in development time and allowed us to select families that differed the most. However, because each female could only lay in one vial per treatment, development time of each family was measured from a single vial. We therefore used experiment 2 to replicate families within and across treatments, and to also measure body size and fitness. Because not all families could be maintained, we chose the 34 fastest and 34 slowest of the families available, and maintained them in vials (c.100 individuals) under standard conditions (25°C, standard diet) for eight months (∼10 generations) before experiment 2.

### Experiment 2: Quantifying selection on development time

We picked eggs for the 68 families into a full-factorial combination of two temperature extremes (14°C and 28°C) and two diets (standard and low-calorie). To pick eggs, we put c.150 flies of each family into two laying pots containing a thin layer of standard food containing 2× the normal amount of agar. To assay productivity (our fitness proxy) at the same time for all treatments, we staggered egg-picks using development times expected from previous experiments, so that flies from all treatments emerged at the same time (**Table S2**). For each family, we picked 20 eggs into each of five vials per treatment (n=340 vials/treatment; total N=1360 vials). We counted all adults that emerged from each vial to estimate egg-to-adult viability. We measured time to eclosion daily for the 14°C treatments, and every six hours for the 28°C treatments.

We measured body size and productivity on one female from each development vial, which means that we measured n=5 females/family/treatment (n=340 females per treatment combination; total N=1360 females). In total, we assayed productivity for 1,242 females, of which 965 had complete data for all traits and were used in the selection analyses. To measure productivity, we collected the fourth female that eclosed from each development vial and placed her into a vial with a randomly selected male from the same treatment. This avoided the earliest-developing females, so that females were closer to the vial mean. We then allowed that female to lay eggs in three vials across a laying period adjusted for each treatment (**Table S2**) to capture the most fecund period while keeping larval density low enough to not affect survival (Flatt 2020). Because females did not lay eggs at 14°C in previous experiments, we assayed productivity at 16°C. Offspring developed at the experimental temperature (with paper cards for pupation), and we counted the adult offspring as our fitness proxy. At the end of the trial, we weighed the wet mass of each focal female using a microbalance (Mettler Toledo, XP2U) and then froze them in individual tubes. We measured wing size as the centroid size of ten wing-vein landmarks on the left wing, photographed under a Leica M80 microscope and quantified using *imageJ* (Walter *et al*. 2026a).

### Statistical analyses

All analyses were conducted in R v.4.5.2 (R Core Team 2025). To test how development time changed across diet and temperature, we used *glmmTMB* (Brooks *et al*. 2017) to apply the linear mixed model

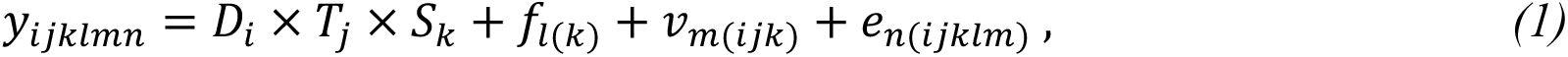

where we tested the interaction between the fixed effect of diet (*D_i_*), temperature (*T*_j_) and chosen families (*S_k_*; fast vs slow developers in experiment 1) on development time (*y_i_*_j*klmn*_). Random effects included replicate family within fast vs slow developers (*f_l_*_(*k*)_) and replicate vial nested within family, diet and temperature (*v_m_*_(*i*j*k*)_). *e_n_*_(*i*j*klm*)_ is the residual. We used *emmeans* (Lenth 2019) to extract the marginal means, and *car* (Fox & Weisberg 2019) to test the significance of interactions with type III ANOVA. To check whether the fast and slow tails chosen in experiment 1 persisted in experiment 2, we also calculated the correlation between family means in experiment 1 and experiment 2 for each treatment.

To quantify phenotypic selection, we used *glmmTMB* to apply generalised linear mixed effects models. We included productivity as the response variable, which we modelled using a zero-inflated negative binomial distribution to account for overdispersion and excess zeros, so that fitness combines the probability of reproducing and the number of offspring produced. We analysed each treatment separately and included body mass, wing size and development time as predictor variables. We assayed productivity in sets of females that emerged at the same time, which we included as a random effect. Predictor traits were standardised to a mean of zero and standard deviation of one. To test for significant non-linear selection, we added the quadratics for each trait as well as the cross-products for each combination of traits. This approach captures both directional selection and curvature in the response surface using traditional selection analyses (Lande & Arnold 1983). We then transformed the coefficients from the log-link scale so that they matched linear and non-linear selection gradients on the scale of relative fitness (Morrissey & Goudie 2022).

To test whether selection changed across treatments, we fitted a single model to all four treatments, with productivity as the response. Fixed effects were temperature, diet and their interaction, together with the linear, quadratic and correlational terms for the three traits (standardised within each treatment), and the interactions of each of these terms with temperature, diet and temperature × diet. As above, we used a zero-inflated negative binomial distribution, included assay set (which was specific to each treatment) as a random effect, and estimated the dispersion and zero-inflation parameters separately for each treatment. Including the non-linear terms substantially improved model fit compared with a model containing only linear terms (ΔAIC = 27). For each trait, we also used likelihood ratio tests to compare models with and without the temperature × diet × trait interaction, which tests whether the effect of diet on linear selection depended on temperature. After removing this three-way interaction, we compared models with and without the diet × trait and temperature × trait interactions to test whether diet or temperature changed linear selection on that trait. We tested for changes in non-linear selection in the same way, by removing the interactions of all six quadratic and correlational terms with temperature × diet, diet, or temperature.

Because traits can only be measured on adults, phenotypic selection estimated from individuals only captures selection on reproductive output among survivors (fecundity selection; Arnold & Wade 1984). To check whether patterns of selection were consistent at early life stages, we used family means to estimate genotypic selection on egg-to-adult viability and total fitness (viability × productivity), and tested whether diet and temperature influenced selection (Rausher 1992; details in **Methods S1**).

To quantify genotype-by-environment interactions in development time, we used *MCMCglmm* (Hadfield 2010) to apply linear mixed models with development time as the Gaussian-distributed response variable. Random effects included family and vial. For family, we specified an unstructured covariance matrix for treatment to estimate a 4×4 matrix, which represents among-family variance in development time for each treatment (temperature-diet combination) along the diagonal, and the covariance between treatments on the off-diagonal. From this matrix, we calculated the genetic correlations between treatments using the *cov2cor* function. For vial, we estimated variance among vials within each treatment to specify vial as the independent level of replication. We used non-informative parameter-expanded priors and 1 million iterations (burn-in 100,000), saving 1,000 samples for each posterior distribution. We ensured that effective sample sizes exceeded 85% of saved samples. For each treatment, we calculated broad-sense heritability (H^2^) as the among-family variance divided by total variance, and also evolvability (I_G_) as the among-family variance divided by the squared treatment mean (Houle 1992).

To estimate plasticity in development time, we used the marginal family means (extracted from the *glmmTMB* models using *emmeans*) and calculated plasticity as the difference between treatments divided by the treatment with the longest development time (Valladares, Sanchez-Gomez & Zavala 2006). This captures plasticity for each genotype as the change in development time relative to the slowest treatment. We calculated plasticity across temperature (for each diet), and then across diets (for each temperature). For each treatment, we then used linear regression of the relative productivity of each family on its plasticity to test the association between plasticity and fitness.

## Results

### Temperature caused larger plastic changes in development time than diet

**Fig. 1a** shows the tails of the distribution in development time chosen from the first experiment. In experiment 2, development time was strongly influenced by temperature and diet, with significant main effects (temperature χ²₁ = 3.37 × 10⁴, *p* < 0.001; diet χ²₁ = 767.0, *p* < 0.001), and a significant temperature × diet interaction (χ²₁ = 30.7, *p* < 0.001). The effect of temperature was substantially larger than that of diet, with a 4–5-fold increase in development time from 28°C to 14°C (both diets), whereas the low-calorie diet created a more modest increase in development time from the standard diet (c.30% at 14°C and c.20% at 28°C; **Fig. 1b**). We only found weak evidence of an overall difference between slow and fast developers (χ²₁ = 3.43, *p* = 0.064), and no evidence that these differences varied across environmental conditions (temperature × diet × selected: χ²₁ = 0.25, *p* = 0.62). Slow-selected families therefore only developed slightly more slowly in all treatments, with no significant difference to the fast-selected families (**Fig. 1b**). Family means in experiment 1 (measured from a single vial per family) were only weakly correlated with those in experiment 2 (r = 0.11–0.30), so the fast and slow tails chosen in experiment 1 were largely not preserved in experiment 2. We therefore combined fast-and slow-selected families in all subsequent analyses.

**Fig. 1.**
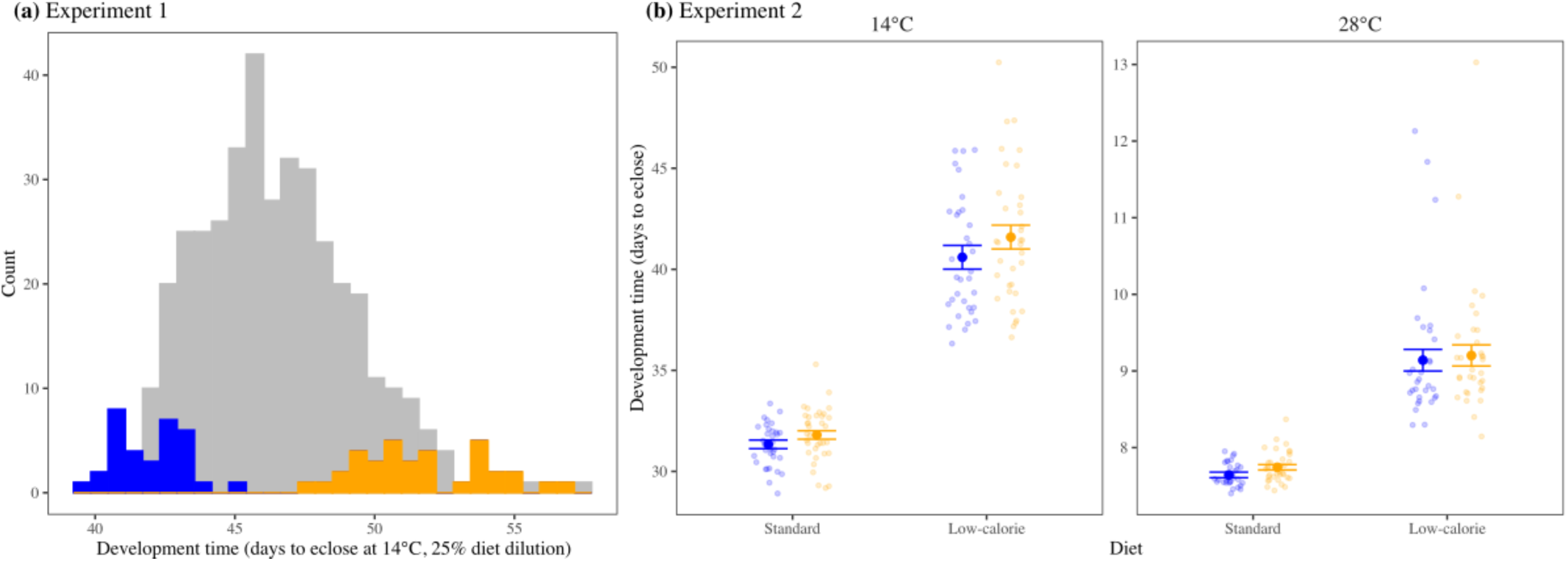
Estimates of development time in both experiments. **(a)** Distribution of development time for the 452 full-sibling families with development time data in experiment 1 under 14°C, low-calorie diet. The chosen fast families are in blue, and slow families in orange. **(b)** Estimates of development time for the fast (blue) and slow (orange) families in each of the four treatments in experiment 2. Small circles represent families, and the large circles with confidence intervals represent mean development time (±1 SE).

### Body size and development time were more strongly associated under nutritional stress

Development time and body size (both wing size and body mass) were weakly correlated when females developed on the standard diet at both temperatures, but moderately and negatively correlated when females developed on the low-calorie diet at both temperatures, such that slower-developing females were smaller (**Fig. 2**). Rearing temperature had little effect on this relationship.

**Fig. 2.**
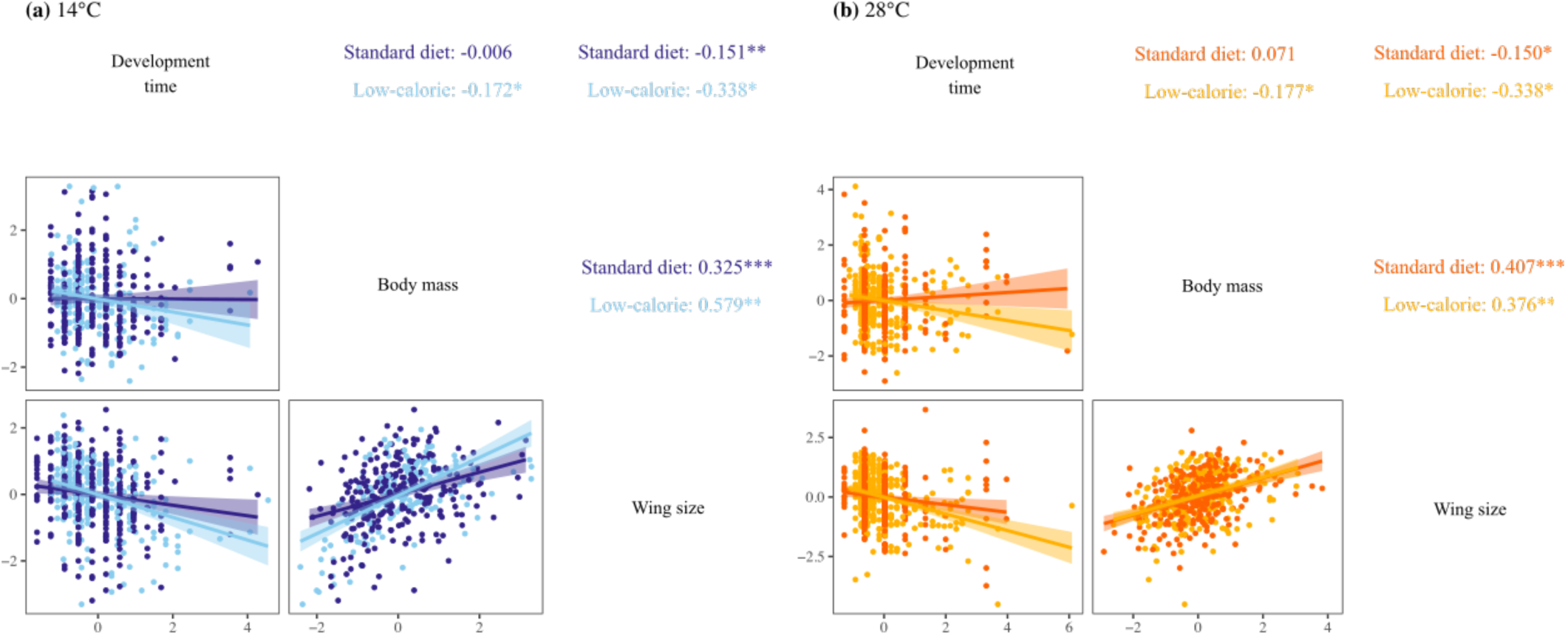
Scatterplot matrices for development time, body mass and wing size estimated for individuals at **(a)** 14°C (blue) and **(b)** 28°C (orange). Darker colours represent the standard diet, and lighter colours the low-calorie diet. Lower diagonals present the scatterplot with a linear regression (±1 SE). Upper diagonal presents the Pearson correlation coefficients for each diet with asterisks denoting significance: *<0.05, **<0.01 and ***<0.001.

### Diet, but not temperature, changed the relationship between development time and fitness

We found no evidence of directional selection on body mass (**Table 1**; **Fig. 3a**). However, directional selection for larger wings was positive in all treatments, and significant on the low-calorie diet at both temperatures (**Table 1**; **Fig. 3b**). Faster development was favoured in the standard diet at 14°C, with a weaker, non-significant trend in the same direction at 28°C, whereas slower development tended to be favoured in the low-calorie diet at 14°C (**Table 1**; **Fig. 3c**). We also found evidence for significant non-linear selection in three treatments (14°C, standard diet: χ²_6_ = 22.57, *p* < 0.001; 28°C, standard diet: χ²_6_ = 38.40, *p* < 0.001; 28°C, low calorie: χ²_6_ = 15.31, *p* = 0.018), which was created by intermediate body mass being favoured at 28°C on both diets, and by correlational selection between development time and body mass at 14°C on the standard diet (**Table 1**; **Fig. 3a**). Apparent non-linear selection on development time under the standard diet at 14°C was driven by two extreme observations, which did not affect directional selection or other estimates when removed.

**Fig. 3.**
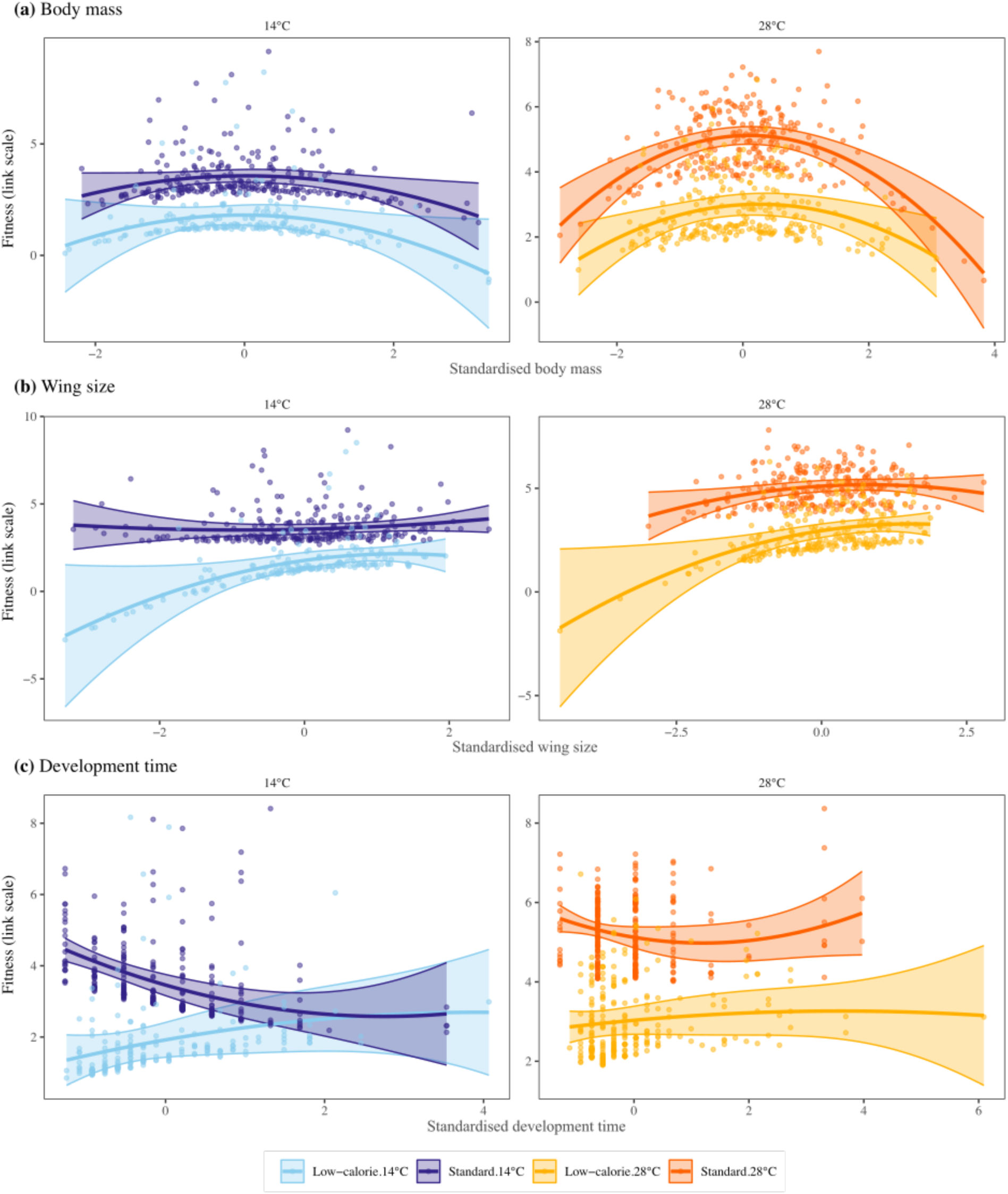
Model-based predictions showing the partial relationships between fitness and **(a)** body mass, **(b)** wing size, and **(c)** development time across nutritional (standard and low-calorie) and thermal (14°C and 28°C) treatments. Lines show fitted relationships from treatment-specific selection models (including non-linear selection terms), and representing the effect of each trait on fitness while accounting for other traits in the model. Shaded regions indicate 95% confidence intervals and points represent partial residuals. Fitness is shown on the link (log) scale of the fitted negative binomial models. Curvature was evident only for body mass, where a concave (hump-shaped) relationship indicates that intermediate body mass was favoured, significantly so at 28°C (**Table 1**).

**Table 1.** Coefficients of selection on development time and body size derived from multiple regression applied to each of the four treatments. Parameters represent the coefficient, with one standard error in parentheses. Parameters in bold are significant at P<0.05, based on Wald tests of the model coefficients on the log-link scale. ***β*** indicate estimates of directional selection. Non-linear selection coefficients include quadratics denoted by grey shading along diagonal, and cross-products below the diagonal.

|  |  |  |  | Non-linear selection coefficients |  |  |
|---|---|---|---|---|---|---|
| Temp. | Diet | Trait | ***β*** | Body mass | Wing size | Development time |
| 14°C | Standard | Body mass | -0.148 (0.119) | **-0.172 (0.154)** |  |  |
|  |  | Wing size | 0.192 (0.145) | -0.108 (0.149) | 0.182 (0.169) |  |
|  |  | Dev. time | **-0.815 (0.237)** | **0.439 (0.216)** | -0.344 (0.25) | 1.079 (0.565) |
|  | Low-calorie | Body mass | 0.158 (0.151) | -0.289 (0.163) |  |  |
|  |  | Wing size | **0.484 (0.213)** | 0.419 (0.222) | -0.07 (0.33) |  |
|  |  | Dev. time | 0.429 (0.215) | 0.308 (0.204) | 0.078 (0.268) | 0.138 (0.169) |
| 28°C | Standard | Body mass | 0.063 (0.055) | **-0.384 (0.059)** |  |  |
|  |  | Wing size | **0.117 (0.061)** | 0.132 (0.069) | -0.144 (0.088) |  |
|  |  | Dev. time | -0.298 (0.201) | -0.107 (0.104) | 0.064 (0.072) | 0.337 (0.273) |
|  | Low-calorie | Body mass | 0.046 (0.065) | **-0.297 (0.072)** |  |  |
|  |  | Wing size | **0.337 (0.078)** | 0.062 (0.072) | -0.108 (0.137) |  |
|  |  | Dev. time | 0.115 (0.13) | -0.018 (0.06) | 0.012 (0.087) | -0.02 (0.054) |

In a single model across all four treatments, diet changed linear selection on development time (diet × development time: χ²₁ = 7.15, p = 0.007), but this effect of diet did not differ between temperatures (temperature × diet × development time: χ²₁ = 0.41, p = 0.52). By contrast, temperature did not change linear selection on development time (temperature × development time: χ²₁ = 1.68, p = 0.20). Diet also strengthened selection for larger wings (diet × wing size: χ²₁ = 8.32, p = 0.004), whereas linear selection on body mass did not change with diet or temperature (all p > 0.30). Non-linear selection changed with temperature (χ²₆ = 18.92, p = 0.004) but not with diet (χ²₆ = 6.59, p = 0.36), with no interactive effects on non-linear selection (temperature × diet: χ²₆ = 3.13, p = 0.79). **Table 1** suggests that temperature effects on non-linear selection were mainly driven by positive correlational selection between development time and body mass at 14°C that was absent at 28°C, and stronger selection for intermediate body mass at 28°C.

We found consistent results for genotypic selection gradients using family means: faster development was associated with higher fitness on the standard diet at both temperatures, and selection on development time was more positive on the low-calorie diet than on the standard diet at both temperatures (**Fig. S1; Table S1**). Including egg-to-adult viability did not change our conclusions (**Results S1; Table S1**). Across individual-and family-level analyses, the low-calorie diet therefore consistently shifted selection towards slower development.

### Nutritional stress increased genetic variation in development time

Genotypic (among-family) variance standardised by the mean (evolvability) was 10–16× greater on the low-calorie diet (I_G_ = 0.65–0.96) than on the standard diet (0.06–0.07), and was similar between temperatures within each diet (**Table 2**). Broad-sense heritability showed the same pattern, increasing from 0.07 on the standard diet at both temperatures to 0.33–0.48 on the low-calorie diet. Genetic correlations well below 1 indicate that families ranked differently across treatments, providing strong evidence of G×E. Correlations between treatments ranged from −0.12 to 0.58 (**Table 2**). In particular, the genetic correlation between diets was −0.12 at 14°C, and 0.23 at 28°C, which suggests that genetic variation in development time was almost independent between diets. Genetic correlations across temperatures were also weaker under nutritional stress (standard diet: 0.58; low-calorie diet: 0.10; **Table 2**).

**Table 2.**
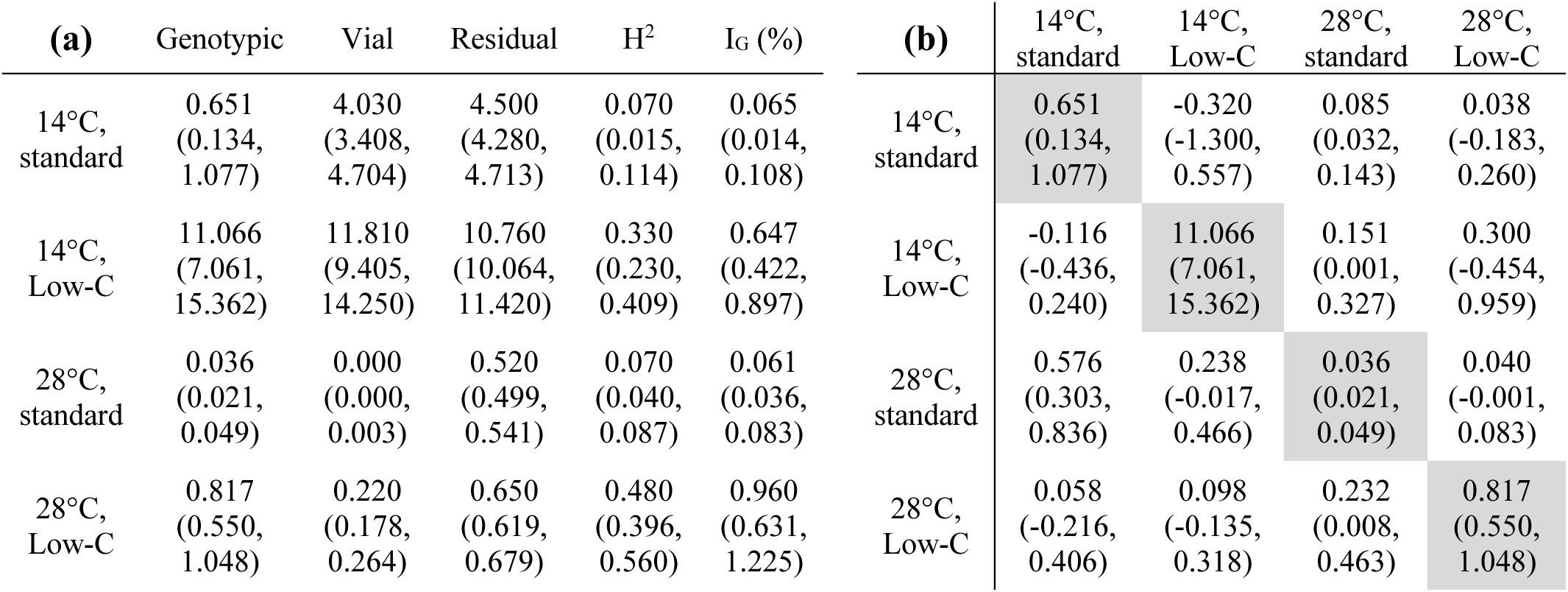
Quantifying G×E interactions for development time across temperature and diet treatments. Presented are the posterior means of the distributions, with the 90% Highest Posterior Density (HPD) intervals in parentheses. **(a)** Partitioning variance among genotypes, vials and residual for each treatment. H^2^ is broad-sense heritability, and I_G_ is evolvability. **(b)** Matrix with genotypic variances along the diagonal (grey shading), genotypic covariances above the diagonal, and genotypic correlations below the diagonal. Correlations near one would suggest that genotypes show the same development time in two given treatments, whereas negative and near-zero correlations suggest that genotypes differ in development time among treatments.

### No evidence that selection favours plasticity in development time

We estimated selection on plasticity as the regression of the relative fitness (productivity) of each family on its plasticity in development time. Slope estimates were near zero and non-significant, and so we found no evidence that plasticity was associated with fitness (**Fig. S2**).

## Discussion

Development time is a functional life history trait that integrates growth, resource acquisition, and reproductive timing (Gotthard, Nylin & Wiklund 1994; Nylin & Gotthard 1998; Flatt 2020), yet we still lack a clear understanding of how its fitness consequences change across realistic combinations of stressors. We tested how temperature and nutritional stress jointly shape the plasticity of development time, its relationship with fitness, and its genetic variation. We show that while temperature created larger plastic changes in mean development time (**Fig. 1b**), nutritional stress shifted selection towards slower development and increased genetic variation in development time (**Tables 1–2**, **Fig. 3**). The stressor that caused the smaller plastic response therefore had the larger effect on directional selection and evolvability. These results suggest that changes in food availability may have a stronger influence than temperature on the adaptive landscape of a key life-history trait, so how populations respond to warming may depend on the nutritional conditions they experience.

### Nutritional stress, but not temperature, changed directional selection on development time

Selection favoured faster development under the standard diet at both temperatures, but this advantage disappeared under the low-calorie diet, where selection shifted towards slower development (**Table 1**). By contrast, temperature did not influence directional selection on development time, indicating that nutritional stress alters the selective landscape for development time. Similarly, in wild common lizards, warmer temperatures strongly advanced the timing of reproduction, whereas directional selection remained constant across 31 years (Bodineau *et al*. 2026). One explanation for our observed shift in selection across diets is that changes in nutrition could determine how development costs arise during life-history (Marshall *et al*. 2020; Abarca *et al*. 2024). When resources are abundant, faster development may be advantageous because it reduces exposure to juvenile mortality and accelerates reproduction (Chippindale *et al*. 1997; Nylin & Gotthard 1998; Prasad & Joshi 2003). Under nutritional stress, however, rapid development may constrain growth or energy accumulation, such that individuals may benefit from prolonging development to accumulate energy reserves or reach physiological maturity (Stearns & Koella 1986; Monaghan 2008). This benefit is unlikely to arise through larger size because slower-developing females were smaller on the low-calorie diet (**Fig. 2**), and selection on development time was estimated while holding body size constant. Physiological studies in *Drosophila* support this interpretation, showing that suboptimal nutrition and temperature reduce lipid and glycogen stores, which constrains energy allocation to reproduction (Klepsatel, Wildridge & Gáliková 2019; Knoblochová *et al*. 2024). Under such conditions, selection may then be strongest on traits that enable better energy storage, such as structural body size, which is consistent with our observation of stronger selection for larger wings under nutritional stress (diet × wing size interaction; **Table 1**). In contrast with our results, limited larval food generated viability selection against long development in yellow dung flies (Blanckenhorn 1998), so the fitness consequences of slower development under nutritional stress may depend on the type of resource limitation.

### Diet mediates strong genotype-by-environment interactions in development time

While temperature had little effect on genetic variation, nutritional stress increased genetic variance and evolvability in development time, and also weakened genetic correlations across diets (**Table 2**). These results suggest that the expression of genetic variation in development time is highly sensitive to nutritional conditions, which may reflect differences in the predictability or consistency of environmental variation experienced by the population (Via & Lande 1985; Ghalambor *et al*. 2007). Temperature variation is common in the wild, whereas diet may vary less predictably (Cross *et al*. 2015; Rosenblatt & Schmitz 2016). Genotypes may therefore be more canalised against changes in temperature than in diet, at least under more benign nutritional conditions (Hoffmann & Sgrò 2011; Chakraborty, Sgrò & Mirth 2020). Nutritional stress can also directly affect developmental and metabolic pathways, which could expose genetic variation in growth and resource allocation (Mirth & Shingleton 2012; Flatt 2020), as is often found for life-history traits under stress (Hoffmann & Merilä 1999; Pallares *et al*. 2023). The weak genetic correlations in development time across diets (**Table 2**) indicate that genotypes that develop quickly on one diet do not necessarily develop quickly on the other. As a consequence, evolutionary responses to selection on development time are likely to be strongly context dependent, with limited potential for correlated responses across environments (Sgrò & Hoffmann 2004).

### Plasticity in development time: variation without fitness benefits

Despite substantial among-family variation in plastic responses of development time across diets and temperatures, we found no evidence that plasticity was associated with fitness (**Fig. S2**). Instead, fitness variation was determined by trait values expressed within each environment, which is consistent with studies showing that variation in plastic responses is often decoupled from mean trait values and may not be directly associated with fitness (Acasuso-Rivero *et al*. 2019; Love & Ferris 2024), or only within a limited range of environments (Żygadło *et al*. 2025). One interpretation is that plasticity in development time reflects physiological constraints, which allow individuals to track environmental variation without plasticity itself being favoured by selection (DeWitt, Sih & Wilson 1998; Nylin & Gotthard 1998). At the level of treatment means, however, plasticity and selection were aligned across diets but not across temperatures (**Fig. 4**). The low-calorie diet both lengthened development and shifted selection towards slower development, whereas temperature caused much larger plastic changes without changing directional selection (**Fig. 4**). Plasticity in response to diet may therefore move the population mean in the direction favoured by selection, even though more plastic genotypes did not have higher fitness.

**Fig. 4.**
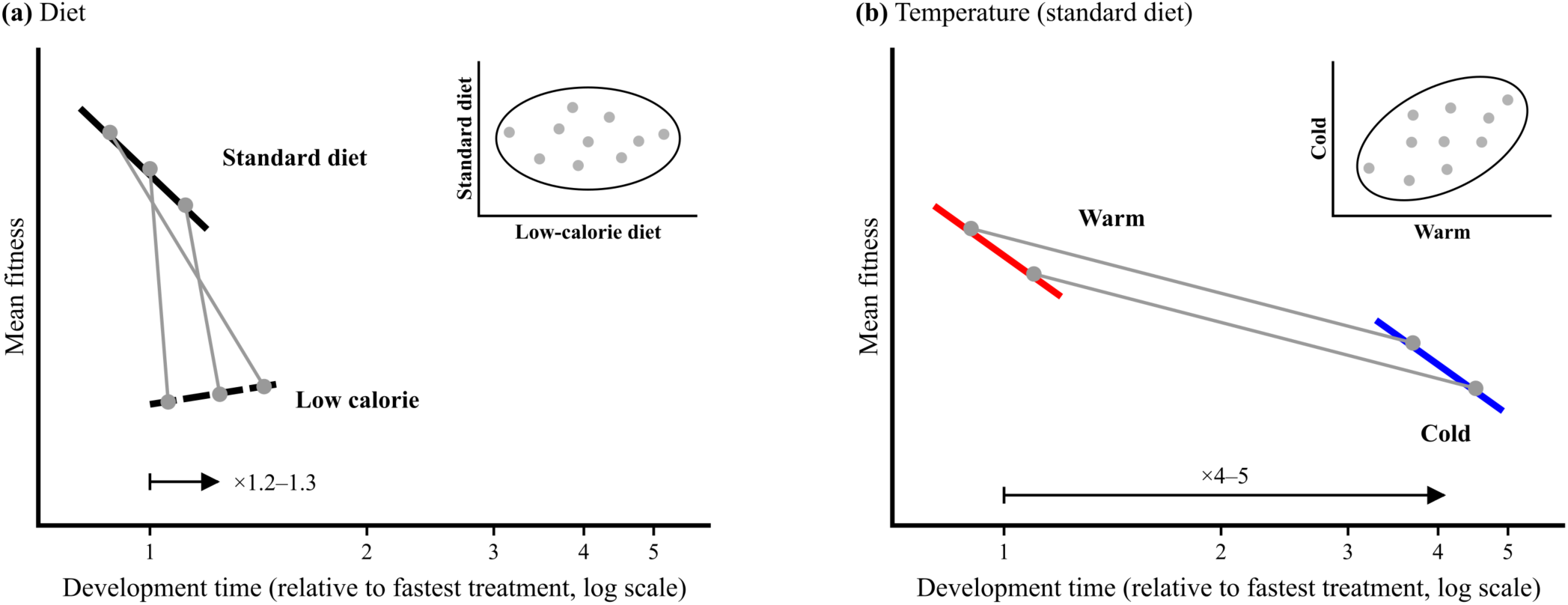
Schematic of the main results across **(a)** diets and **(b)** temperatures. Lines show the direction of selection, and grey lines connect representative genotypes across environments. The x-axis shows development time relative to the fastest treatment (log scale), and arrows show the mean plastic change (diet: ×1.2–1.3; temperature: ×4–5). Insets plot genotypes (grey circles) in the two environments, with ellipses showing the genetic correlation. **(a)** Nutritional stress slowed development and shifted selection from faster towards slower development, matching the direction of plasticity. Genotypes partly changed rank, so genetic correlations across diets were close to zero (inset; **Table 2**). **(b)** Temperature caused much larger plastic changes without changing directional selection, and genetic correlations across temperatures were positive on the standard diet (inset) but close to zero on the low-calorie diet (**Table 2**).

### Implications for adaptive changes in development time under multiple stressors

Together with previous work, our results suggest that temperature and nutrition act at different scales. Temperature largely determined the strength of latitudinal clines in stress tolerance and body size (Walter *et al*. 2025), and within our population it caused much larger plastic changes in development time than diet (**Fig. 1b)**. By contrast, diet had a larger effect on directional selection and genetic variation within the population, increasing genetic variance in development time and creating near-zero genetic correlation in development time across diets (**Table 2**). Therefore, although we predicted that temperature and diet would interact, we found that their effects on development time were largely independent. The only exception was that nutritional stress weakened the genetic correlation in development time across temperatures (standard diet: 0.58; low-calorie diet: 0.10; **Table 2**). By contrast, in the same population, temperature and diet stress interacted to increase genetic variance in fitness, and while the strength of genetic correlations changed across combinations of diet and temperature, they remained positive and moderately strong (Walter *et al*. 2026b). Although estimated on different sets of families, these contrasting results suggest that genotypes that perform well on one diet tend to perform well on the other, but achieve this with different development times, which is consistent with diet changing how development time influences fitness. Because food availability does not vary consistently with latitude, adaptation to nutrition may be local and leave little signature across broad-scale clinal populations (Chakraborty, Sgrò & Mirth 2020; Walter *et al*. 2025). While temperature appears to structure broad-scale patterns of adaptation, nutritional stress may play a more prominent role in shaping contemporary selection and the availability of genetic variation within populations (Chakraborty *et al*. 2023; Choy *et al*. 2024). Predicting life-history responses to global change will therefore require understanding how plasticity, selection and evolutionary potential of functional traits respond to multiple environmental axes (Gienapp *et al*. 2008; Chevin, Lande & Mace 2010; Urban *et al*. 2024).

### Caveats

Because our population combined flies from across a 300×600 km area, some G×E may reflect historical adaptation to different environments, even though high gene flow should mean that these genes segregate across the broader population. Fitness was measured as productivity over a defined reproductive window, and viability could only be included at the family level, which means that we could only measure a proxy for lifetime reproductive success. Our estimates of broad-sense genetic variance includes non-additive effects (Sgrò & Blows 2003; Rudin-Bitterli, Mitchell & Evans 2018) that may not translate into evolutionary responses (Lynch & Walsh 1998), and may explain why heritability of development time did not change across similar diets when estimated from additive variance (Gebhardt & Stearns 1992).

Families were also maintained for c.10 generations before experiment 2, and inbreeding depression could have increased among-family variance on the low-calorie diet. However, broad-sense heritability for development time in experiment 1 (14°C, low-calorie) was 0.26 (0.23–0.29 90% HPD), which is similar to that of experiment 2 in the same treatment (H² = 0.33; 0.23–0.41), so inbreeding is unlikely to explain the 10–16-fold increase in evolvability under nutritional stress. In addition, nutritional stress also increased additive genetic variance in fitness in this outbred design (Walter *et al*. 2026b), consistent with development time in the current study. These limitations are therefore unlikely to explain our main result that nutritional stress shifted selection towards slower development and increased genetic variance.

## Conclusions

Our results show that the fitness consequences of development time depend more on nutrition than on temperature, even though temperature induced larger plastic changes in development time. While faster development was favoured under benign nutritional conditions, this advantage disappeared when food was limited, and selection shifted towards slower development. These shifts were accompanied by strong genotype-by-environment interactions that increased the expression of genetic variance under nutritional stress, highlighting how environmental stress reshapes both phenotypic optima and evolutionary potential. More generally, the size of the plastic response in development time was a poor guide to how the environment changed its contribution to fitness, suggesting that the functional significance of plastic traits needs to be assessed from the relationship with fitness (i.e., measuring selection and genetic variation) within each environment. To predict evolutionary responses to global change, a key next step will be to test whether patterns of selection and genetic variation observed here translate into realised evolutionary responses under combinations of stressors.

## Author contributions

GW designed the experiments, collected and analysed the data, and wrote the paper. AC and FC collected the data. MH helped to analyse the data and guided the framing of the paper. LA, CW and CS provided facilities, experimental support, and helped with experimental design. All authors contributed to the writing.

## Acknowledgements

We are very grateful to Isobel Tyson and Connor Bevan for their help conducting the second experiment, and we thank Emily Lombardi and Lina Vo for their help with picking eggs. GMW is grateful to have been supported by Australian Research Council fellowships DE200101019 and FT240100466. CMS was supported by Australian Research Council Discovery Projects DP220102958 and DP240102830. CRW and LAA were supported by Australian Research Council Discovery Projects DP180103925, DP220103553 and DP22010342, and LAA was also supported by an Australian Research Council fellowship FT250100816.

