## Supplementary material for "Why development time matters: temperature influences plasticity while nutritional stress reshapes selection and evolutionary potential"

**Contents**

**Fig S1** Genotypic selection gradients.

**Fig S2** Selection gradients on plasticity.

**Methods S1 and Results S1** Selection through viability and total fitness.

**Table S1** Genotypic selection gradients for productivity, viability and total fitness.

**Table S2** Expected development times and productivity assay schedule in each treatment.

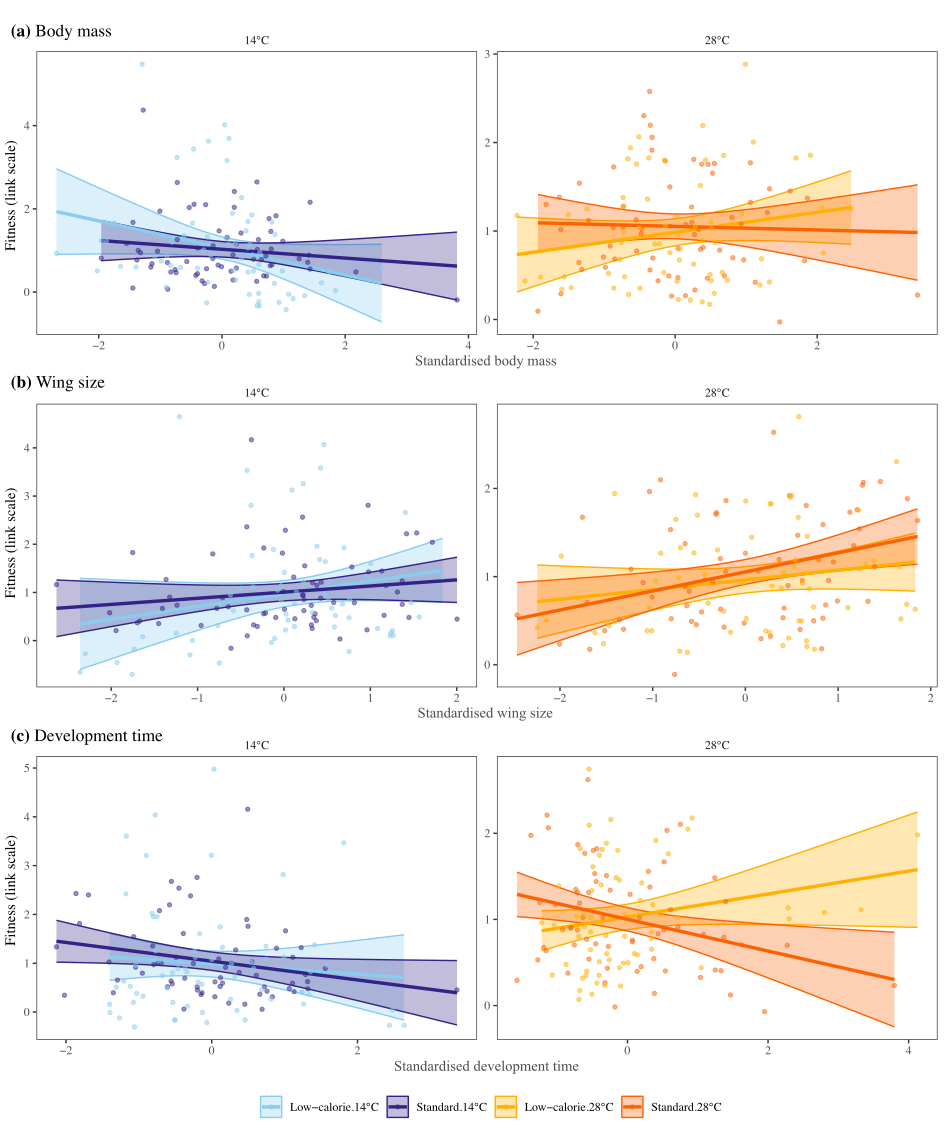

**Fig S1** Genotypic selection gradients. Each point represents the marginal mean for a full-sibling family. Fitness is relative productivity, calculated from family marginal means estimated using a negative binomial mixed model. Only linear selection has been estimated because we found no evidence of non-linear selection on genotypic means.

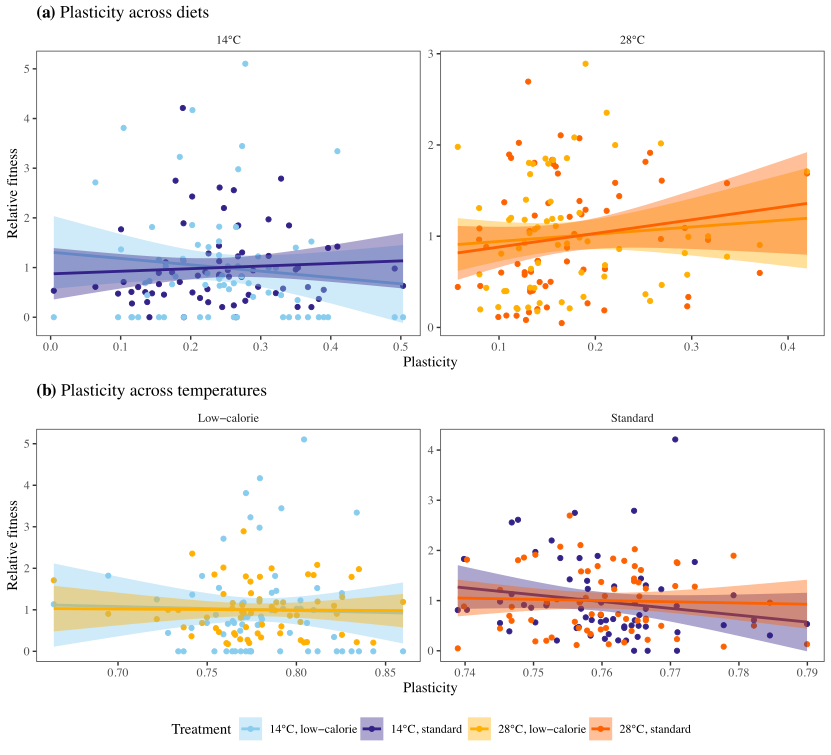

**Fig S2** Selection on plasticity across **(a)** diets, and **(b)** across temperatures. We found no evidence for selection on plasticity.

**Methods S1 Selection through viability and total fitness.**

For each treatment, we estimated family-mean egg-to-adult viability from the number of adults that emerged from the 20 eggs in each vial, using a beta-binomial model (*glmmTMB*) with family as a fixed effect, and extracted the marginal family means using *emmeans*. We removed two vials that recorded more than 20 adults, which were errors during egg-picking. We then calculated total fitness for each family as the product of its viability and its mean productivity, which represents the expected number of adult offspring produced per egg. We estimated linear genotypic selection gradients for each fitness component by regressing relative fitness on the family means for body mass, wing size and development time, standardised within each treatment. We tested whether diet or temperature altered genotypic selection on development time using a single linear model across all four treatments, which included temperature, diet and their interactions with each trait. We used F-tests to compare models with and without the temperature × diet × development time interaction and, after removing it, with and without the two-way interactions (diet × development time and temperature × development time interactions).

**Results S1 Selection through viability and total fitness.** Egg-to-adult viability was lowest at 14°C on the low-calorie diet (mean 0.26, compared with 0.40–0.47 in the other treatments). In all four treatments, viability selection on development time had the same sign as productivity selection. Using total fitness, faster-developing families had higher fitness on the standard diet at both temperatures, but not on the low-calorie diet (**Table S1**). In a single model across all four treatments, diet altered genotypic selection on development time through total fitness (diet × development time: F_1,252_ = 6.41, p = 0.012) and viability (F_1,252_ = 7.16, p = 0.008), with a marginal effect for productivity (F_1,252_ = 3.70, p = 0.056). These effects of diet did not differ between temperatures (all p > 0.28). Temperature altered selection on development time through viability (temperature × development time: F_1,252_ = 4.32, p = 0.039), but not through productivity or total fitness (both p > 0.15). The individual-level selection for slower development at 14°C on the low-calorie diet (**Table 1**) was not seen among families, where selection on development time was close to zero for all fitness components. Overall, including viability supported our conclusion from the analysis of productivity in the main text: the low-calorie diet shifted selection on development time towards slower development at both temperatures, whereas temperature altered selection only through viability and not through total fitness.

**Table S1** Genotypic selection gradients (β, with one standard error in parentheses) for body mass, wing size and development time, estimated from family means for three fitness components: productivity (number of adult offspring per surviving female), egg-to-adult viability, and total fitness (viability × productivity, i.e. the expected number of adult offspring per egg). Each fitness component is relative to the treatment mean, and traits were standardised within each treatment. Values in bold are significant (p < 0.05).

| **Temp.** | **Diet** | **Fitness component** | **Body mass** | **Wing size** | **Development time** |
| --- | --- | --- | --- | --- | --- |
| 14°C | Standard | Productivity | −0.106 (0.113) | 0.128 (0.113) | −0.190 (0.096) |
| 14°C | Standard | Viability | −0.014 (0.056) | **0.124 (0.056)** | **−0.150 (0.047)** |
| 14°C | Standard | Total fitness | −0.214 (0.161) | 0.254 (0.161) | **−0.284 (0.137)** |
| 14°C | Low-calorie | Productivity | −0.326 (0.188) | 0.263 (0.189) | −0.104 (0.161) |
| 14°C | Low-calorie | Viability | −0.029 (0.078) | **0.223 (0.078)** | −0.042 (0.067) |
| 14°C | Low-calorie | Total fitness | −0.267 (0.176) | 0.350 (0.177) | −0.103 (0.151) |
| 28°C | Standard | Productivity | −0.020 (0.080) | **0.216 (0.081)** | **−0.184 (0.074)** |
| 28°C | Standard | Viability | −0.040 (0.044) | **0.108 (0.045)** | −0.070 (0.041) |
| 28°C | Standard | Total fitness | −0.045 (0.099) | **0.290 (0.102)** | **−0.226 (0.092)** |
| 28°C | Low-calorie | Productivity | 0.113 (0.088) | 0.110 (0.088) | 0.133 (0.082) |
| 28°C | Low-calorie | Viability | 0.034 (0.050) | 0.082 (0.050) | 0.091 (0.047) |
| 28°C | Low-calorie | Total fitness | 0.140 (0.114) | 0.198 (0.115) | 0.211 (0.107) |

**Table S2** Expected development times and productivity assay schedule in each treatment of experiment 2. Expected development times, from previous experiments, were used to stagger egg-picks so that flies from all treatments emerged at the same time. Females did not mature or lay eggs at 14°C, so females from the 14°C treatments were assayed for productivity at 16°C. Each female laid eggs in three vials spread equally across the laying period, which was adjusted for each treatment to capture the most fecund period while keeping larval density low.

| **Treatment** | **Expected development time (days)** | **Eclosion scored** | **Productivity assay temperature** | **Laying period (days)** |
| --- | --- | --- | --- | --- |
| 14°C, standard | 38 | Daily | 16°C | 18 |
| 14°C, low-calorie | 44 | Daily | 16°C | 22 |
| 28°C, standard | 8 | Every 6 h | 28°C | 9 |
| 28°C, low-calorie | 10 | Every 6 h | 28°C | 12 |
